# Capturing multiple timescales of adaptation to second-order statistics with generalized linear models: gain scaling and fractional differentiation

**DOI:** 10.1101/2019.12.30.891143

**Authors:** Kenneth W. Latimer, Adrienne L. Fairhall

## Abstract

Single neurons can dynamically change the gain of their spiking responses to account for shifts in stimulus variance. Moreover, gain adaptation can occur across multiple timescales. Here, we examine the ability of a simple statistical model of spike trains, the generalized linear model (GLM), to account for these adaptive effects. The GLM describes spiking as a Poisson process whose rate depends on a linear combination of the stimulus and recent spike history. The GLM successfully replicates gain scaling observed in Hodgkin-Huxley simulations of cortical neurons that occurs when the ratio of spike-generating potassium and sodium conductances approaches one. Gain scaling in the GLM depends on the length and shape of the spike history filter. Additionally, the GLM captures adaptation that occurs over multiple timescales as a fractional derivative of the stimulus variance, which has been observed in neurons that include long timescale afterhyperpolarization conductances. Fractional differentiation in GLMs requires long spike history that span several seconds. Together, these results demonstrate that the GLM provides a tractable statistical approach for examining single-neuron adaptive computations in response to changes in stimulus variance.

## 1 Introduction

Neurons adapt their spiking responses in a number of ways to the statistics of their inputs (Fairhall, 2014). A particularly well-studied example is adaptation to the stimulus variance, which can provide important computational properties. First, neurons can show gain scaling, such that the input is scaled by the stimulus standard deviation (Fairhall et al., 2001a; Mease et al., 2013). Scaling of the gain by the stimulus standard deviation implies that single spikes maintain the same information about the stimulus independent of its overall amplitude. This adaptation of the “input gain” with stimulus standard deviation can occur very rapidly. Second, the mean firing rate can adapt to variations in the stimulus variance across multiple timescales (Fairhall et al., 2001b; Wark et al., 2007). This form of spike frequency adaptation can in some cases have power-law properties (Pozzorini et al., 2013) and serve to compute the fractional derivative of the variance (Anastasio, 1998; Lundstrom et al., 2008).

One approach to studying such adaptation is to use Hodgkin-Huxley style (HH) conductance based models to explore potential single-neuron mechanisms underlying these computations (Lundstrom et al., 2008; Mease et al., 2013). Although HH models can indeed capture such behavior, the mechanistic HH framework is not ideally suited for statistical analysis of spike train data in sensory systems as HH model parameters are difficult to interpret in terms of computation and coding. Moreover, fitting HH models to intracellular data is difficult (Buhry et al., 2011; Csercsik et al., 2012; Vavoulis et al., 2012; Lankarany et al., 2014), and only recently methods that fit HH models to spike trains alone have been gaining success (Meng et al., 2011, 2014).

In contrast, statistical point process models based on the generalized linear model (GLM) framework have provided a tractable tool for modeling spiking responses of neurons in sensory systems (Truccolo et al., 2005; Pillow et al., 2008). Previous work has shown the utility of finding linear features that can explain the spiking behavior of HH models (Agüera y Arcas et al., 2003; Agüera y Arcas and Fairhall, 2003; Weber and Pillow, 2017). Unlike simple linear/nonlinear models, GLMs also incorporate a dependence on the history of activity, potentially providing a helpful interpretative framework for adaptation (Mease et al., 2014). We therefore fit GLMs to spike trains generated from a range of HH neurons. We found that the GLMs could reproduce the single-neuron adaptive computations of gain scaling and fractional differentiation. Capturing gain scaling across a range of HH active conductance parameters depended both on the choice of link function and spike history length. As the length of the spike history filter increased, the stimulus dependency of neurons changed from differentiating to integrating (Stevenson, 2018). Capturing adaptation as a fractional derivative required a history fflter that could account for long timescale effects: on the order of 10 s. Together these results demonstrate that the GLM provides a tractable statistical framework for modeling adaptation that occurs at the single-neuron level.

## 2 Materials and Methods

### 2.1 Gain scaling

Gain scaling refers to the case when for an input-output function of a neuron, the input gain is proportional to the standard deviation (SD) of the stimulus (*σ*). Thus, the gain depends on the recent context. If a neuron achieves perfect gain scaling, the firing rate *R* given a particular stimulus value, *s*, and input standard deviation can be written as:

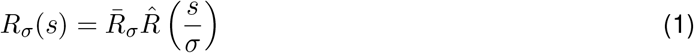

where the normalized stimulus 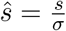, and the output gain, 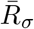, is constant in *s*.

To quantify the degree of gain scaling in a neuron’s spiking output, we measure the firing rate function in response to a white-noise input, *x*(*t*), at different SDs and constant mean *μ* (**Figure 1A**). For each standard deviation, we compute the normalized spike-triggered average (STA; **Figure 1B**) (Rieke et al., 1999). We then compute the stimulus as the convolution 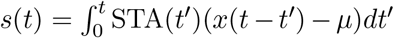. The spike rate function is then defined probabilistically as

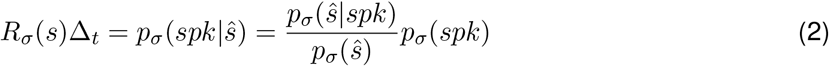

where the right side follows from Bayes’ rule. The average firing rate in time bin of width Δ_*t*_ is *p_σ_*(*spk*). Thus, we get 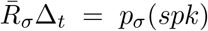 and 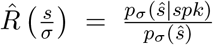. The spike-triggered stimulus distribution, *p_σ_*(*ŝ|spk*), is the probability of the stimulus given that a spike occurred in the bin. By definition the marginal stimulus distribution, *p_σ_*(*ŝ*), is a standard normal distribution which does not depend on *σ*. Therefore, if *p_σ_*(*ŝ|spk*) is similar across different values of *σ*, gain scaling is achieved because 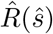 does not depend on *σ*.

**Figure 1:**
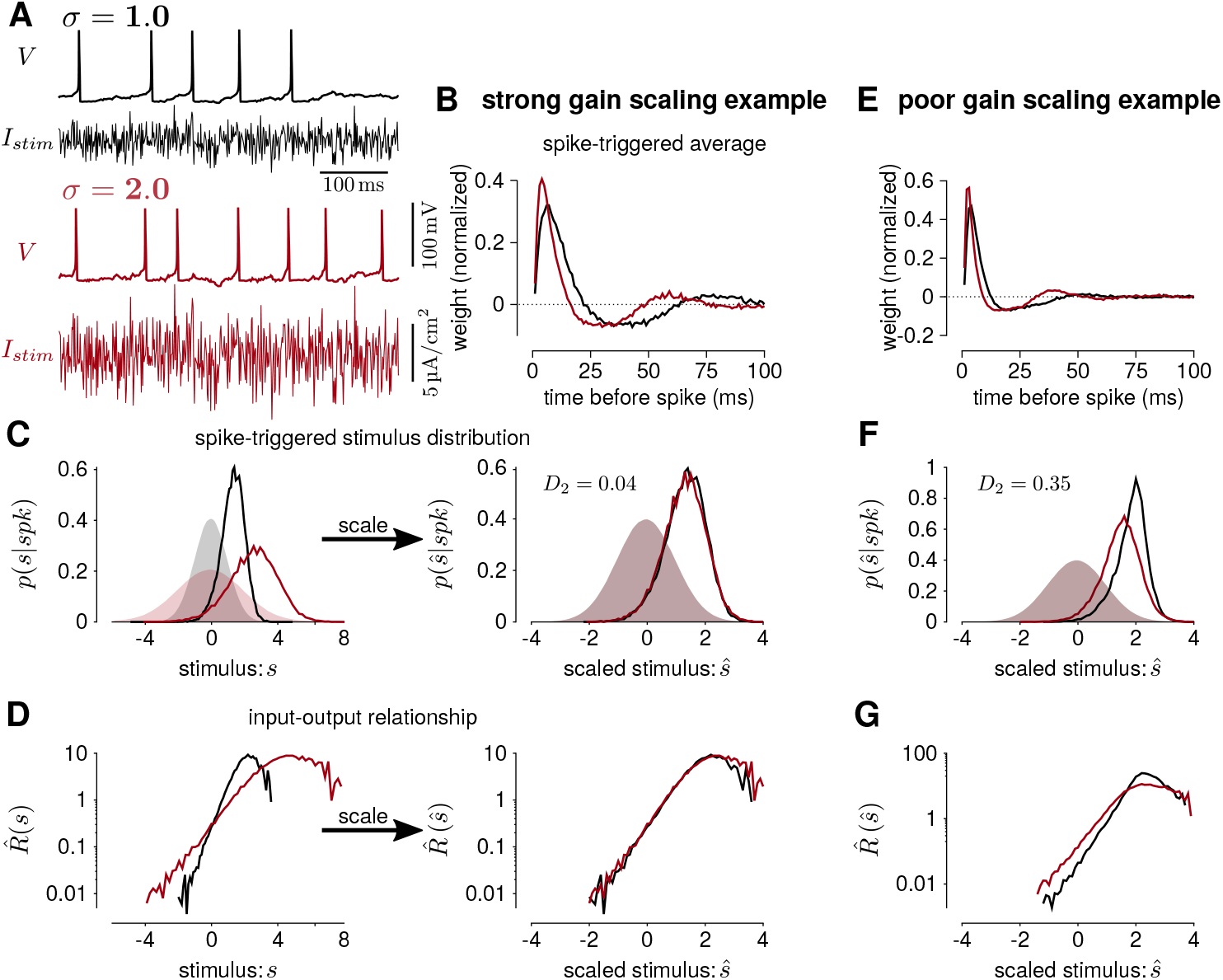
(**A**) Hodgkin-Huxley simulation of a neuron stimulated with white noise at two different standard deviation levels (black *σ* = 1; red *σ* = 2). In this simulation, the total sodium and potassium conductances were equal (*G_Na_* = *G_K_* = 1000 pS/μm^2^). (**B**) The STAs measured at the two stimulus standard deviations. (**C**) Left shows the spike-triggered distributions of the STA filtered input (*s*) and right shows the distributions over the STA filtered input scaled by the standard deviation (*ŝ*). The shaded areas show the prior stimulus distributions, which are Gaussian distributed with standard deviation *σ*. (**D**) The input-output functions of the stimulation at each stimulus level. Scaling the input by the standard deviation shows that the simulated neuron scales the gain of the input by the stimulus standard deviation (right). (**E**) The STAs measured at two standard deviations from a Hodgkin-Huxley simulation with high potassium and low sodium total conductances (*G_Na_* = 600 and *G_K_* = 2000 pS/μm^2^). The spike-triggered stimulus distribution (**F**) and scaled input-output function (**G**) for this simulation does not show gain scaling.

We measure gain scaling in terms of the spike-triggered distribution. We do so using the 1st Wasserstein, or earth-mover’s metric (we obtained qualitatively similar results using the symmetrized Kullback-Leibler divergence and Jensen-Shannon divergence). The Wasserstein metric is a distance function between two probability distributions. Intuitively, it can be thought of as the minimum work needed to transform one distribution into the other by moving probability mass as if the distributions are piles of sand (**Supplementary Figure 1**). Formally, it is defined as

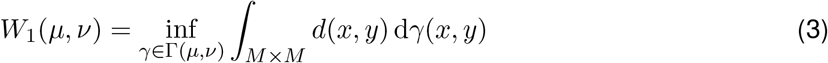

where *ν* and *μ* are probability measures on a metric space *M* with metric *d*(·,·). The infimum is taken over the collection of measures, Γ(*μ*, *ν*), on *M* × *M* with *μ* and *ν* marginal distributions. We compute the gain scaling score at *σ* as *D_σ_* = *W*_1_(*p*_1_(*ŝ*|*spk*), *p_σ_*(*ŝ*|*spk*)). A distance close to 0 indicates that the spike-triggered distributions are similar, and therefore the cell is gain scaling its input (**Figure 1C-D**). Larger values of *D_σ_* indicate that the input-output function does not scale with *σ* (**Figure 1F-G**). We computed the spike-triggered distribution using a histogram with bins of width 0.1.

#### 2.1.1 Gain scaling in Hodgkin-Huxley neurons

A previous study by Mease et al. (2013) found that Hodgkin-Huxley models could account for gain scaling observed in pyramidal neurons. Thus we simulated spikes from single-compartment Hodgkin-Huxley style models of pyramidal neurons, providing a source of data with which to explore the expression of this property using GLMs. The voltage and gating dynamics followed the equations (Mainen et al., 1995)

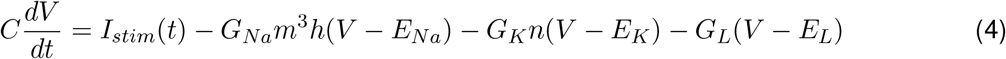

such that for each gate *x* ∈ {*n, m, h*}

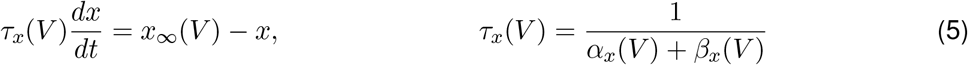

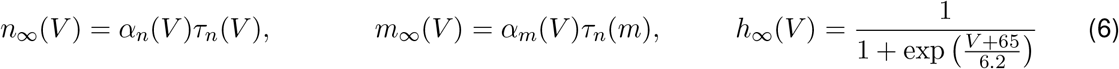

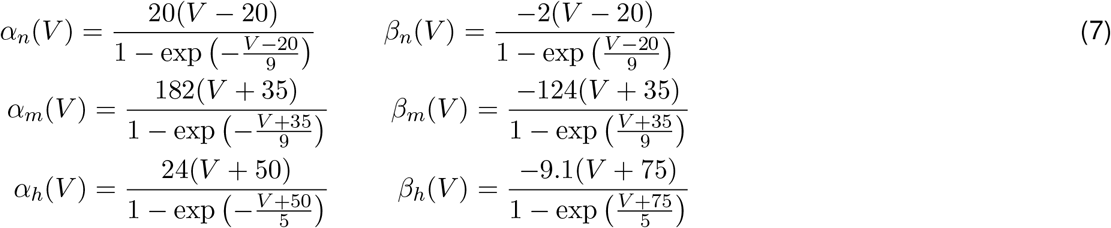

The reversal potentials were *E_Na_* = −70, *E_K_* = 50, and *E_L_* = −70 mV and the capacitance was *C* = 1 μF/cm^2^. The leak conductance was set to 0.4 pS/μm^2^ so that the resting membrane had a time constant of approximately 25 ms. As in Mease et al. (2013), we explored a range of values for the active conductances *G_Na_* and *G_K_*: from 600-2000pS/μm^2^ in increments of 100 pS/μm^2^. Simulations were performed in MATLAB using a fourth-order Runge-Kutta method with step size 0.01 ms. Spike times were defined as upward crossings of the voltage trace at −10 mV separated by at least 2 ms.

The input consisted of Gaussian draws every 1 ms with parameters 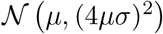 where *σ* was set to 1.0, 1.3, 1.6 or 2.0. For each value of *G_Na_* and *G_K_*, the mean input, *μ*, was tuned so that at baseline, where *σ* = 1, each simulation produced approximately 10 spk/s using a 100 s simulation. We did not consider values of *G_Na_* and *G_K_* that spiked spontaneously (i.e., spiked when *μ* = 0). We simulated 2000 s of spiking activity at each stimulus level (generating approximately 20000 spikes at *σ* = 1).

### 2.2 Fractional differentiation

We next looked at periodic modulations of the stimulus standard deviation to model long timescale adaptive effects. We applied stimuli consisting of Gaussian noise with sinusoidal or square wave modulation of the variance between 1 and *σ* with *σ* again taking values of 1.3, 1.6 or 2.0, at a number of different frequencies. We analyzed simulated spike trains across 7 noise modulations periods: 1,2, 4,8, 16, 32, and 64 s. The simulations were 3200 s for each period, giving a minimum of 50 cycles per period.

Lundstrom et al. (2008) found that pyramidal neurons can act as fractional differentiators of the stimulus amplitude envelope for this type of input. Fractional derivatives generalize the derivative operator such that, analogous to taking the first derivative of a function twice to obtain the second derivative, taking the fractional derivative of order *α* = 1/2 twice results in the first derivative (Oldham and Spanier, 1974). Fractional differential filters respond to a square stimulus as an exponential-like decay with a time constant that depends on α (**Figure 2A-B**). Fractionally differentiating a sinusoidal stimulus produces a *frequency dependent* gain change (**Figure 2C**)

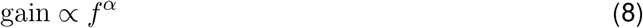

where *f* is the frequency. Additionally, fractionally differentiating the sine function gives a frequency independent phase shift, *ϕ*, of the stimulus (**Figure 2D**):

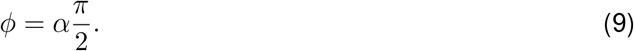

**Figure 2:**
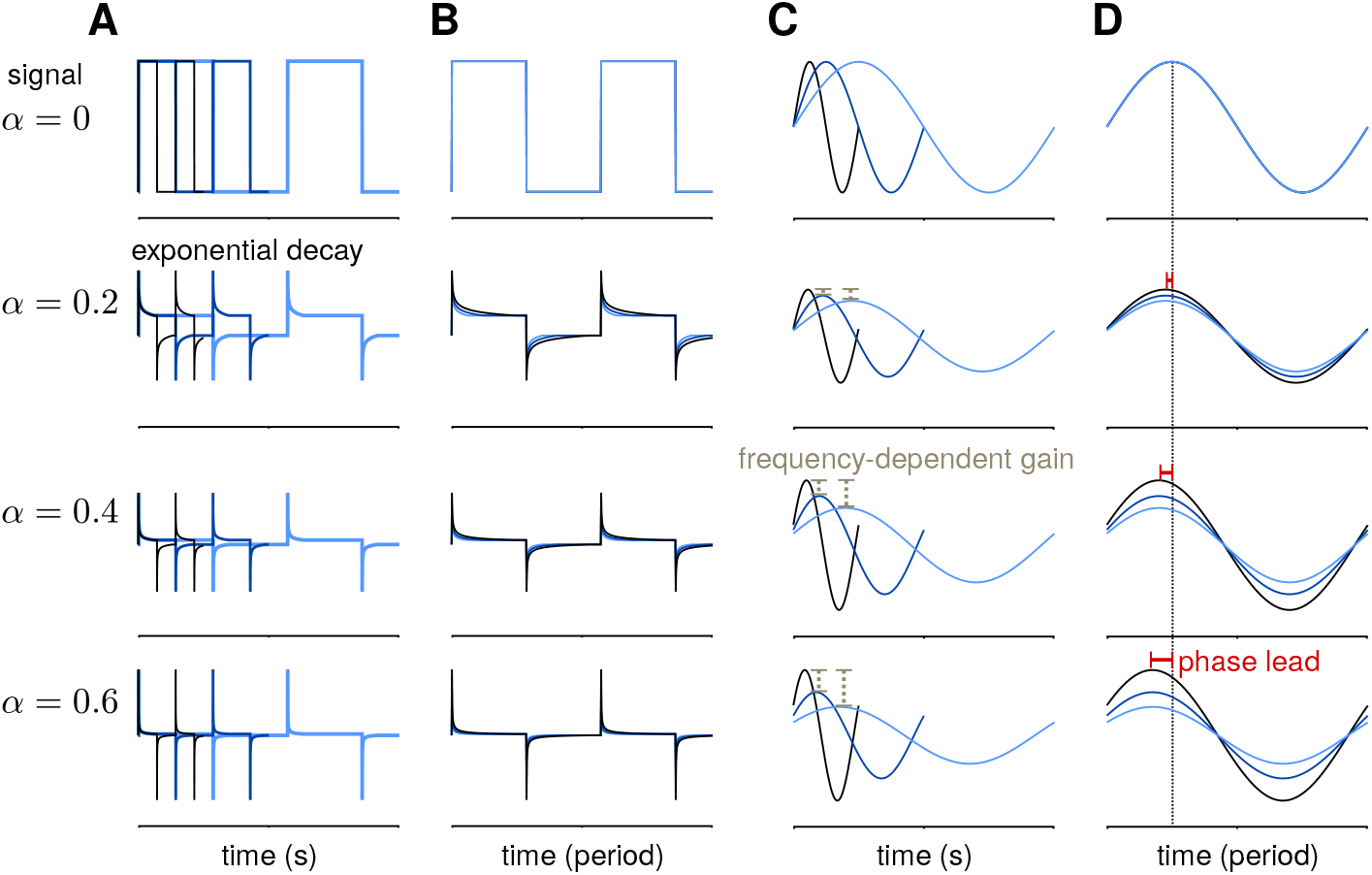
Example of a fractional derivative of several orders. Each row shows a different fractional order (*α*) of the function in the top row. (**A**) The fractional derivatives of a step function with three different periods (colors) shows exponential filtering with an *α* dependent timescale. (**B**) The fractional derivatives in A scaled by period. (**C**) The fractional derivatives of a sine function for three different periods. As *α* increases, the fractional derivative shows greater frequency-dependent gain. (**D**) The same function as in **B** with the sine functions scaled by period. At higher orders, the phase lead of the fractional derivative relative to the signal increases equally over frequencies.

These three measures can be combined to estimate approximate fractional differentiation by neurons.

To compute the fractional derivative order, we computed cycle-averaged responses obtained using 30 bins per cycle at each stimulus amplitude modulation frequency. We fit the cycle-averaged square-wave responses across all modulation frequencies as the best fitting fractional derivative of the stimulus amplitude (plus a baseline rate) using least-squares. To fit *α* to the phase lead of the sine-wave responses, we computed mean phase lead (*ϕ*) across frequencies and applied Equation 9. To fit *α* to the gain of the sine-wave responses, we applied Equation 8 by fitting a least-squares regression between the frequency of modulation and the logarithm of the gain.

#### 2.2.1 Fractional differentiation by Hodgkin-Huxley neurons

We simulated neurons from the standard HH model with three additional afterhyperpolarization (AHP) currents with time constants ranging from 0.3 to 6 s. The equations for the HH neurons were

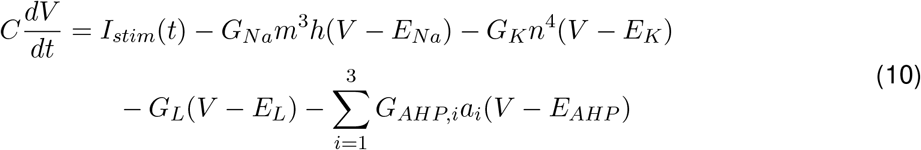

The gates *x* ∈ *n,m,h* follow the dynamics

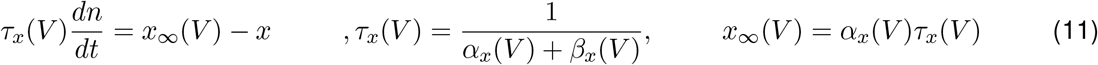

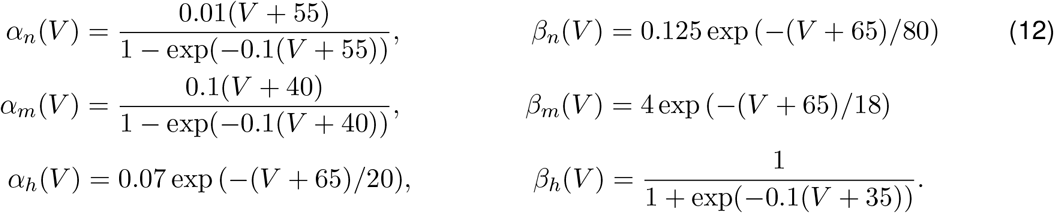

The AHP currents have linear dynamics and are incremented by 1 at spike times (*t_spk,i_*):

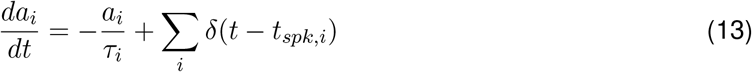

where *δ* is the Dirac delta function. The standard were: *G_Na_* = 120, *G_K_* = 36, *G_L_* = 0.3 mS/cm^2^; *E_Na_* = 50, *E_K_* = −77, *E_l_* = −54.4 mV; and *C* = 1 μF/cm^2^. The AHP conductances were set relative to the leak conductance: *G_AHP_*,. = (0.05, 0.006 and 0.004)*G_L_*. The AHP reversal potential was *E_AHP_* = −100 mV and the AHP timescales were set to *τ_i_* = (0.3, 1, and 6)s.

Similarly to the gain scaling simulations, the stimulus was sampled independently in each 1 ms bin from a normal distribution with mean *μ*. The time-dependent variance given *σ* and the period (*p*) was 4*μf_p_*(*t, σ*). The time-dependent modulation function for the square-wave stimulus was

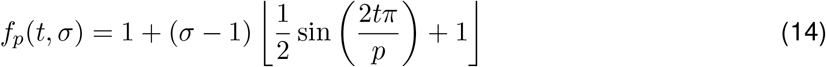

where ⌊·⌋ denotes the floor operator, and the function for the sine-wave stimulus was similarly defined as

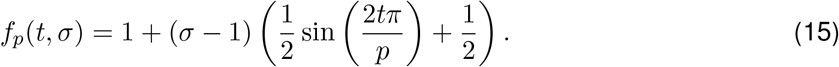

The parameter *μ* was calibrated so that with no variance modulation (i.e., *σ* = 1), the simulated cells produced approximately 10 spk/s.

### 2.3 Generalized linear models

The GLM models the spiking process as an autoregressive Poisson process with (**Figure 3A**). The spike rate at time *t* is given as a linear-nonlinear function of the stimulus and the spike history

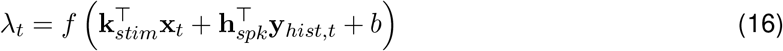

where **x**_*t*_ is the stimulus vector preceding time *t*, and **y**_*hist*_ is the spike history vector. The parameters of the GLM are the stimulus filter (**k**_*stim*_), the spike history filter (**h**_*s*_ *pk*), and baseline rate (*b*). For the inverse-link function, *f*, we used the canonical exponential function except where otherwise noted.

**Figure 3:**
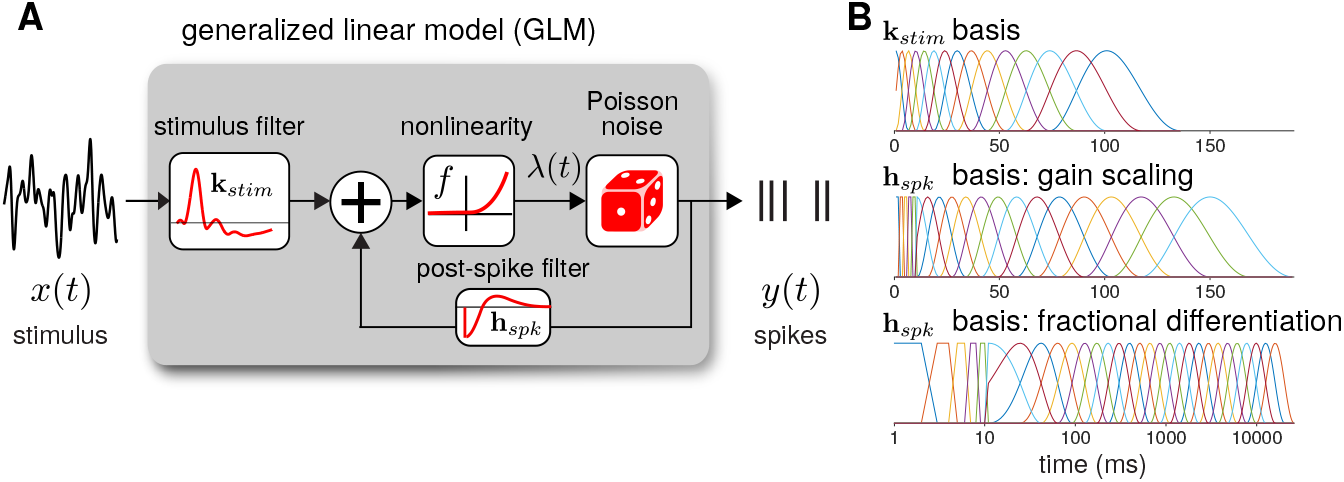
(**A**) Diagram of the neural GLM that describes spiking as an autoregressive Poisson process. (**B**) The basis functions used to parameterize the GLM filters. (top) The stimulus basis used for all GLMs. (middle) The spike history basis used for the gain scaling simulations. (bottom) The spike history basis used for the fractional differentiation simulations. Due to the length of the spike history filters needed to capture fractional differentiation, the time axis is shown in log scale.

The log-likelihood of a binned spike train, **y**, given the model parameters is then

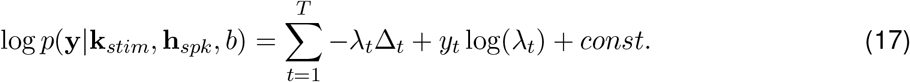

For all model fits and simulations, we set Δ_*t*_ = 1 ms. We numerically maximized the log-likelihood using conjugate-gradient methods.

To reduce the number of model parameters, we parameterized the *S* filters using smooth basis functions (**Figure 3B**). The stimulus filter was parameterized using 15 raised cosine basis functions:

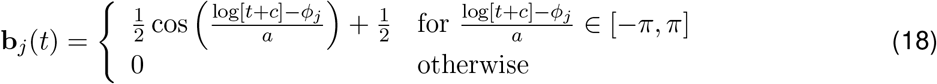

where *t* is in seconds. We set *c* = 0.02 and *a* = 2(*ϕ*_2_ − *ϕ*_1_)/*π*. The *ϕ_j_* were evenly spaced from *ϕ*_1_ = log(*T*_0_/1000 + *c*), *ϕ*_15_ =log(*T_end_*/1000 + *c*) where the peaks of the filters are in the range *T*_0_ = 0 and *T_end_* = 100 ms.

The spike history filter bases were constructed in two parts. To account for the absolute refractory period, we used 5 box car filters of width 2 ms for the first 10 ms of the spike history. The remaining spike history filter was parameterized using raised cosine basis functions with the parameter *c* = 0.05. For the gain scaling simulations, *N* = 15 cosine basis functions were used with spacing *T*_0_ = 10 and *T_end_* = 150 ms. For the fractional differentiation simulations, *N* = 25 cosine basis functions were used with spacing *T*_0_ = 10 and *T_end_* = 16000 ms. To explore how the timescale of spike history affected adaptation in the GLM, for each model we fit the GLM using only the first *i* cosine basis functions for each *i* = 0 (using only the refractory box-car functions) to *i* = *N*. Thus, we obtained *N* + 1 nested model fits across a range of spike history lengths. When stated, the length of the spike history filter, *T_hist_*, denotes the time of the peak of the *i*th basis function.

#### 2.3.1 Evaluating model performance

We evaluated the GLM performance by assessing the ability of the GLM to predict the HH model response to a 32 s novel stimulus. For the gain scaling simulations, we tested the response to the test stimulus at each stimulus SD (*σ*). For the fractional differentiation simulations, the stimulus SD was modulated by a sine or square wave with a 4 s period and a modulation height of *σ* = 2.0. Predictive performance was evaluated using the pseudo-*R*^2^ score (Cameron and Windmeijer, 1997). We selected this measure because it can be applied to Poisson process observations instead of trial-averaged firing rates as is required by the standard *R*^2^ measureof explained variance (Benjamin et al., 2018). Thus, it is especially appropriate for comparing the stochastic GLM to a spike train simulated by the deterministic HH model. The pseudo-*R*^2^ is written as the ratio of deviances:

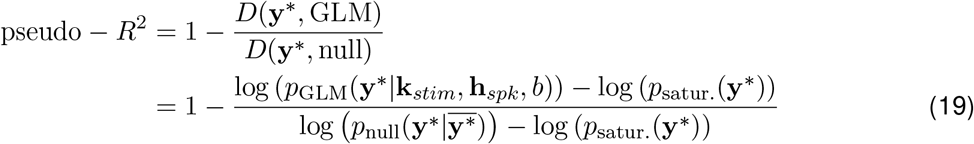

where **y**\* is the test spike train. The GLM likelihood is *p*_GLM_(*Y***y**\*|**k**_*stim*_, **h**_*spk*_, *b*) and the likelihood of the null model 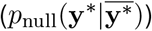 is the probability of the spike train given only the mean firing rate, 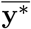. The saturated model likelihood (*p*_satur._(**y**\*)) is the probability of observing **y**\* given one parameter per bin: that is, the Poisson probability observing **y**\* given a model with rate *λ* = 1 in each bin in which the HH model spiked and rate *λ* = 0 in each bin that the HH did not spike. Thus, the pseudo-*R*^2^ measures the fraction of explainable log-likelihood captured by the GLM.

## 3 Results

### 3.1 GLMs capture gain scaling behavior

To investigate how GLMs can capture biophysically realistic gain scaling, we fit the Hodgkin-Huxley simulations with GLMs (**Figure 4A**). We fit a unique GLM for each value of *G_Na_* and *G_K_* in the HH model, and the GLMs were fit using the entire range of stimulus SDs (*σ* = 1.0, 1.3, 1.6, and 2.0). Applying the STA analysis at the four stimulus SDs, we quantified gain scaling in GLM fits and compared the gain scaling in the GLM simulations to the HH neurons (**Figure 4B-C**). Across the range of spiking conductance values, we found that the GLM fits consistently showed gain scaling (**Figure 4D**). The HH neurons showed the greatest degree of gain scaling when the *G_Na_*/*G_K_* ratio was close to one, with the lowest *D*_2_ score occurring at a ratio of 1.17 (Mease et al., 2013). We observed the same pattern in the GLM simulations, but the GLM fits generally exhibited stronger gain scaling when *G_Na_*/*G_K_* < 1 than the HH neurons.

**Figure 4:**
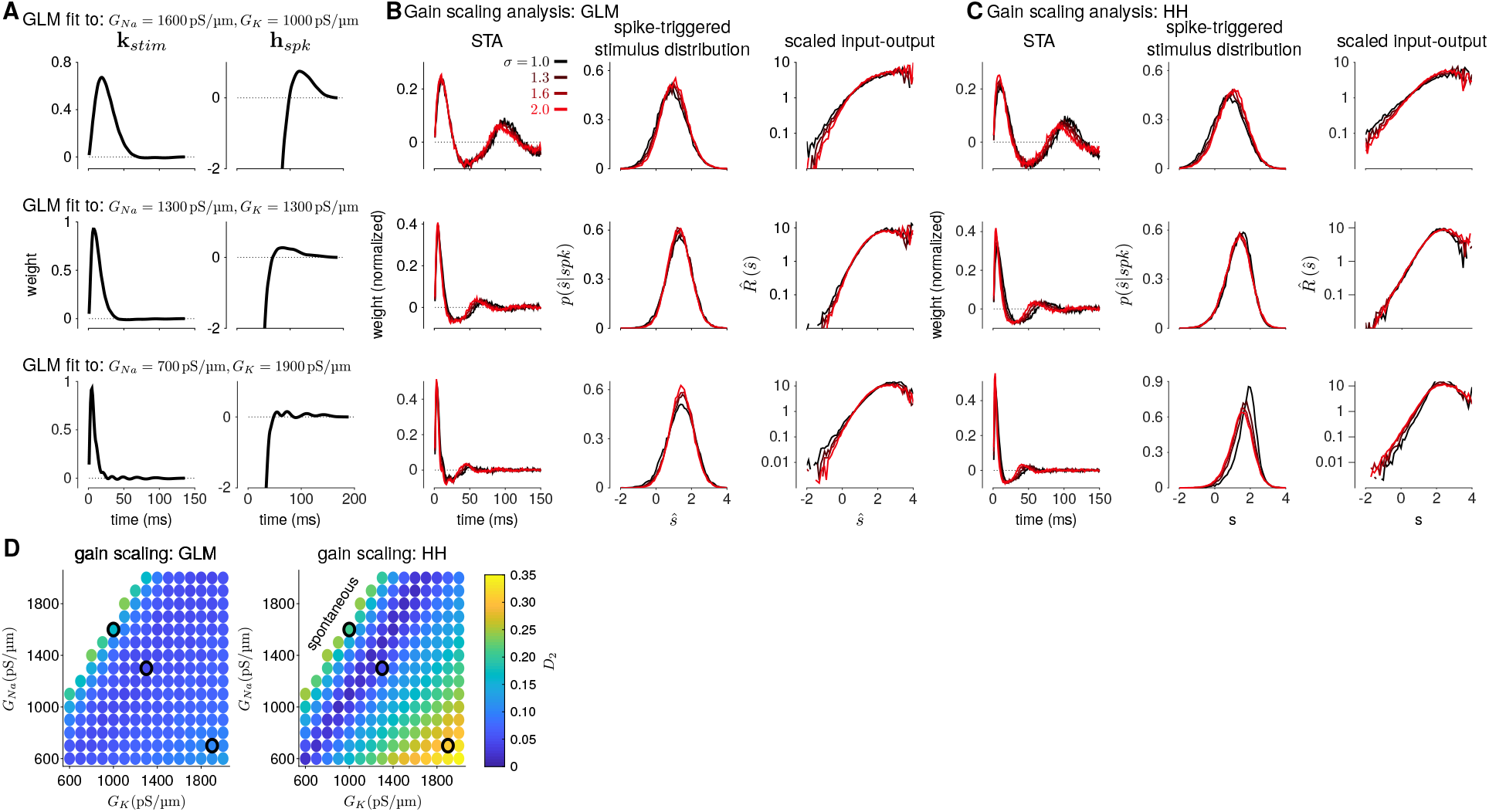
(**A**) Example filters from GLM fits to HH simulations with three different spiking conductance levels (rows). Large negative values driving the refractory period in the spike history filter (right) have been truncated. (**B**) The spike-triggered averages (left), scaled spike-triggered stimulus distributions (right), and scaled input-output functions (right) for the GLM fits in A for all four stimulus SDs. (**C**) Same as B for the HH simulations. (**D**) Gain scaling performance (Wasserstein distance between the spike-triggered distributions at *σ* = 1 and *σ* = 2) at all the spiking conductance levels explored for the GLM simulations (top) and the Hodgkin-Huxley simulations (bottom). Lower values of *D*_2_ correspond to stronger gain scaling. The three black circles indicate the conductance levels for the GLM examples in A and B. Gain scaling was not computed for values of *G_Na_* and *G_K_* that resulted in spontaneous spiking in the Hodgkin-Huxley simulations.

The GLM’s characterization of the HH neurons depended on the spike history filter. This is revealed by comparing the stimulus filters (**Figure 4A**) to the stimulus features extracted by spike-triggered averaging (**Figure 4B**): While the STA showed multiphasic responses, the GLM stimulus filter was consistent with a simple, monophasic integration. This demonstrates that the STA reflects the combination of stimulus and spike history effects (Stevenson, 2018; Agüera y Arcas and Fairhall, 2003). A spike history filter of sufficient length was necessary to achieve accurate model fits across all stimulus SDs (**Figure 5A,B**).

**Figure 5:**
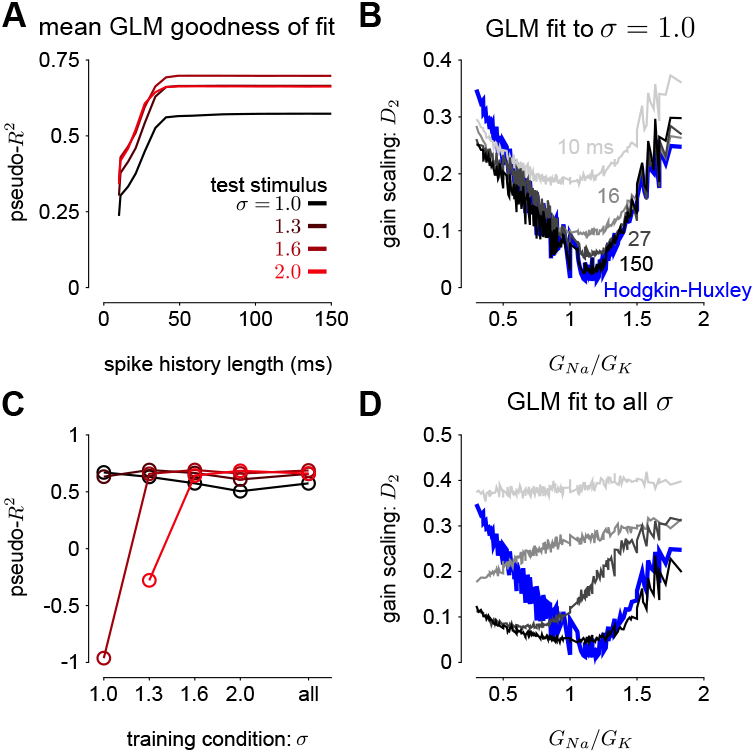
(**A**) The pseudo-*R*^2^ of the GLM fits at the four different stimulus SDs averaged over all *G_Na_* and *G_K_* as a function of spike history length. The GLMs were trained at all stimulus conditions. (**B**) Gain scaling in the Hodgkin-Huxley simulations (blue) measured as a function of the sodium-potassium conductance ratio (*G_Na_*/*G_K_*). The gray traces show gain scaling measured in the GLMs fit to the HH simulations for four different spike history filter length. The GLMs were trained using HH simulations with the stimulus at the baseline SD (*σ* = 1.0). (**C**) The average pseudo-*R*^2^ measured for each *σ* for the GLMs given each training stimulus condition. (**D**) Same as B for the GLMs fit only using all four values*σ*.

We also explored how the stimulus conditions used to fit the GLM determined the model’s ability to capture gain scaling. Remarkably, we found that the GLM fit only to the baseline stimulus SD (*σ* = 1.0) captured the gain scaling pattern seen in the HH neuron (**Figure 5B**). The gain scaling observed in the GLMs required a sufficiently long spike history filter, on the order of at least 50 ms. With shorter spike history, the GLM did not obtain the same level of gain scaling performance at the optimal *G_Na_*/*G_K_* ratio.

However, these GLM fits failed to generalize across stimulus SDs. The GLM trained only at *σ* = 1.0 explained less variance in the spiking responses to a stimulus at *σ* = 2.0 than a model capturing only the mean firing rate for all values of *G_Na_* and *G_K_* (predictive pseudo-*R*^2^ less than 0; **Figure 5C**). Therefore, the GLM trained at *σ* = 1.0 does not accurately characterize the HH responses despite accurately predicting gain scaling in those cells. In contrast, GLMs trained at all four *σ* values failed to capture the lack of gain scaling at low *G_Na_*/*G_K_* values despite showing improved model fit across all *σ* (**Figure 5D**; a detailed example is provided in **Supplementary Figure 2A**). Because the GLM trained on all *σ* showed both consistent generalization performance and strong gain scaling behavior, the remaining analyses considered only that training condition.

We next considered how the GLM parameters related to the gain scaling computation and the space of *G_Na_* and *G_K_* in the HH models. To visualize the geometry of the model parameters, we performed PCA on the stimulus and spike history filters (**Figure 6A,E**). The filters produced across the two HH parameters spanned a two-dimensional subspace (variance explained: stimulus 98.8%, spike history 97.3%). The PCA reconstructions for example stimulus filters are given in **Supplementary Figure 3**. However, the PCs do not correspond to a linear mapping of the *G_Na_* and *G_K_* axes (**Figure 6B,F**). Instead, the first component for both filters correlated with the *G_Na_*/*G_K_* ratio (**Figure 6C,G**; stimulus PC1 *r* = −0.97, *p* < 10^−4^; spike history PC2 *r* = 0.97, p < 10^−4^). The second correlates with the gain scaling value observed in the corresponding HH model (**Figure 6D,H**; stimulus PC2 *r* = −0.89, *p* < 10^−4^; spike history PC2 *r* = 0.90, *p* < 10^−4^). Thus, the GLM parameterizes the HH neuron in a space that corresponds to the ratio *G_Na_*/*G_K_* and gain scaling factor.

**Figure 6:**
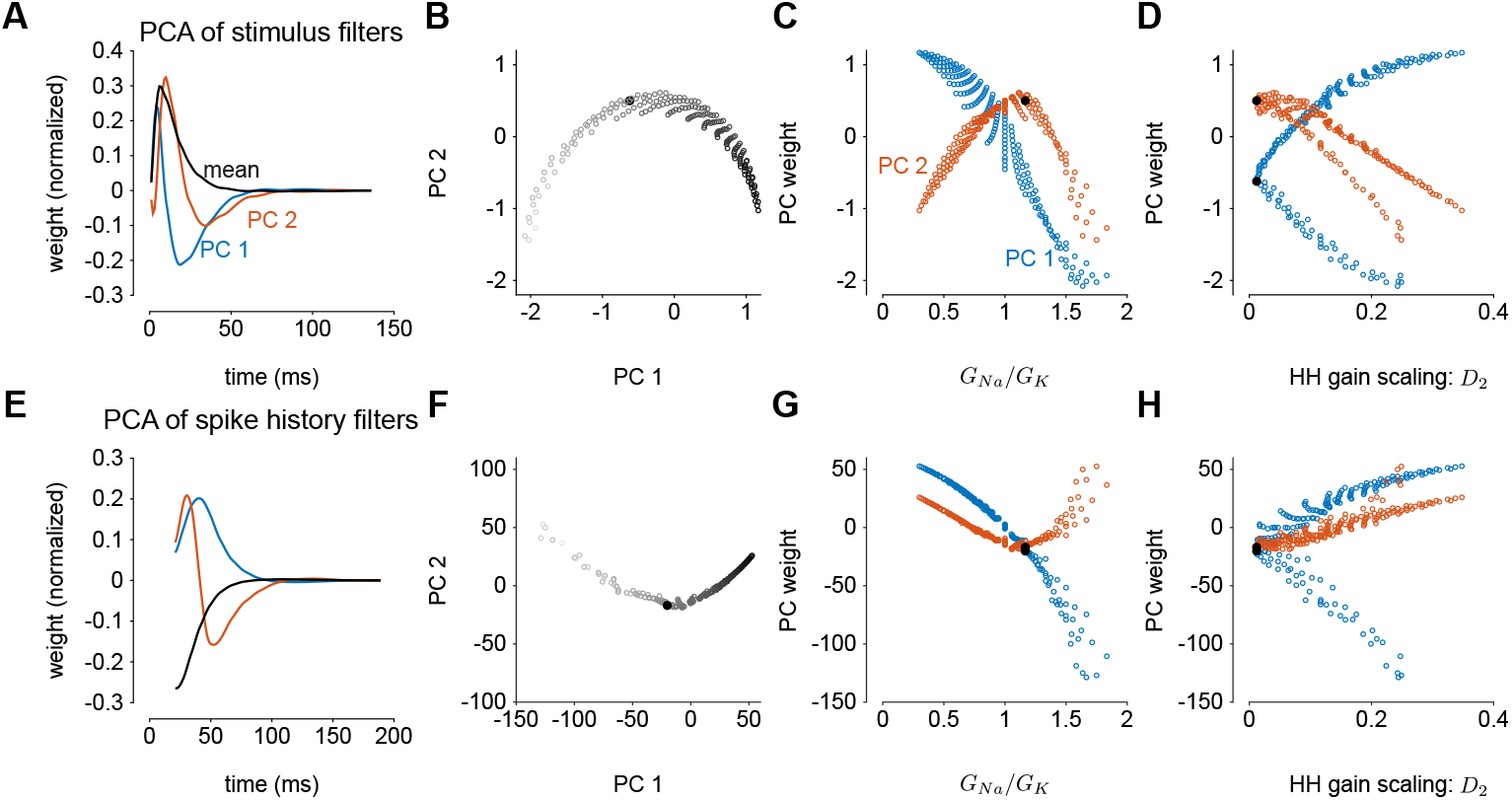
Principal component analysis of the GLM stimulus and spike history filters trained across all values of *G_Na_* and *G_K_*. The GLMs were trained on all *σ* values with a spike history length of 150 ms. (**A**) The first two PCs (blue and red) of the stimulus filter. The normalized mean filter is given in black. (**B**) The projections of the stimulus filters onto the first two PCs. The shade of the points corresponds to *G_Na_*/*G_K_* where lighter indicates a higher ratio. The black points in B-C,E-F indicates GLM fit to the HH model with the best gain scaling (i.e., lowest *D*_2_). (**C**) The stimulus filter PC weights (same as in B) as a function of the *G_Na_*/*G_K_* ratio. (**D**) The stimulus filter PC weights as a function of the gain scaling factor (*D*_2_) observed in the HH simulation fit by the GLM. (**D-F**) Same as A-F for the GLMs’ spike history filters. The first 20 ms of the spike history filters were excluded from analysis to avoid effects from the strong refractory period.

#### 3.1.1 Power-law firing rate nonlinearities

The GLMs we considered used the canonical inverse-link function, the exponential nonlinearity (McCullagh and Nelder, 1989), to transform the filtered stimulus plus spike history into a firing rate. However, it is known that firing rate nonlinearities that instead have a power-law relationship of the input produce gain scaling (Miller and Troyer, 2002; Murphy and Miller, 2003). We therefore considered a range of soft-power nonlinearities over a range of exponents for the GLM firing rate (**Figure 7A**; Equation 16):

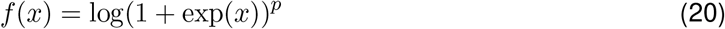

for *p* ∈ {2, 3, 4, 5} (for *p* = 1, the model performed poorly for all HH simulations and the results are not shown). We found that the power-law nonlinearity produced better predictive fit than the exponential for HH simulations with low *G_Na_*/*G_K_* ratios (**Figure 7B**). For those ratios, the exponential GLM in fact predicted *greater* gain scaling than the HH simulation actually showed (**Figure 5A** and **Supplementary Figure 2A**). We found the power-law nonlinearities showed *less* gain scaling in the low *G_Na_*/*G_K_* regime, which was more consistent with the HH simulations (**Figure 7C**). This perhaps counter-intuitive result is likely due to the temporal processing of the GLM: the spike history filter shapes the effective stimulus-response function over longer timescales. Thus, the instantaneous spike rate function need not be a power law to produce gain scaling and an instantaneous power-law function may not result in strong gain scaling in the presence of spike history dependencies.

**Figure 7:**
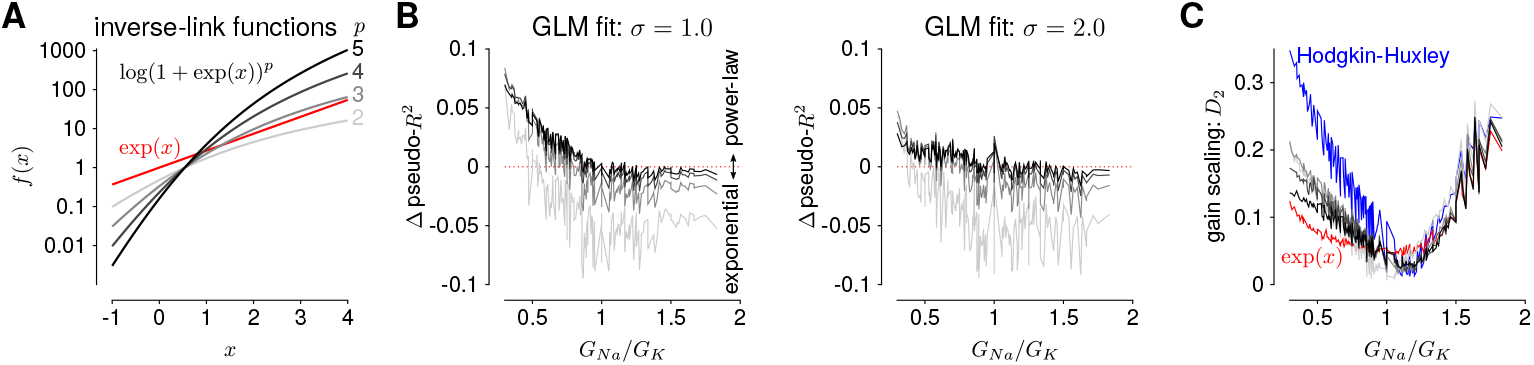
(**A**) The 5 inverse-link functions tested in the GLM. The red trace shows the canonical exponential inverse-link function used in **Figures 4-5**. The gray traces show the soft-power function for different exponents, *p*. (**B**) The difference in predictive performance (measured as pseudo-*R*^2^) between the exponential GLMs and the power-law GLMs for *p* ∈ {2, 3, 4, 5} for a test stimulus of *σ* = 1.0 (left) and *σ* = 2.0 (right). Positive values indicate the GLM with a power-law nonlinearity had greater predictive performance than the exponential GLM. The GLMs were fit to all *σ*. (**C**) Gain scaling predicted by the power-law GLMs (gray traces) compared to the exponential GLMs (red) and the HH simulations (blue).

### 3.2 GLMs capture fractional differentiation with long timescales of adaptation

In this section, we address adaptive computations occurring over multiple timescales spanning tens of seconds, instead of instantaneous gain. We consider adaptation to changes in stimulus variance in the responses of HH simulations with three AHP currents (Lundstrom et al., 2008). The neurons were injected with noise stimuli such with a periodically modulated SD. The stimulus SD followed either a sine or square wave. We focused our analyses on the cycle-averaged firing rate to see how the neural responses reflect fractional differentiation of the stimulus SD envelope in the cycle-averages.

We fit GLMs to HH simulations in response to with either sine- or square-wave SD modulation. The training data included simulations with noise modulation periods of 1 to 64 s. We considered GLMs with different lengths of spike history filters. Cycle-averaged responses of HH and GLM simulations appear qualitatively similar (**Figure 8**), and thus we aimed to characterize how well the GLM fits captured the fractional differentiation properties of the HH neuron.

**Figure 8:**
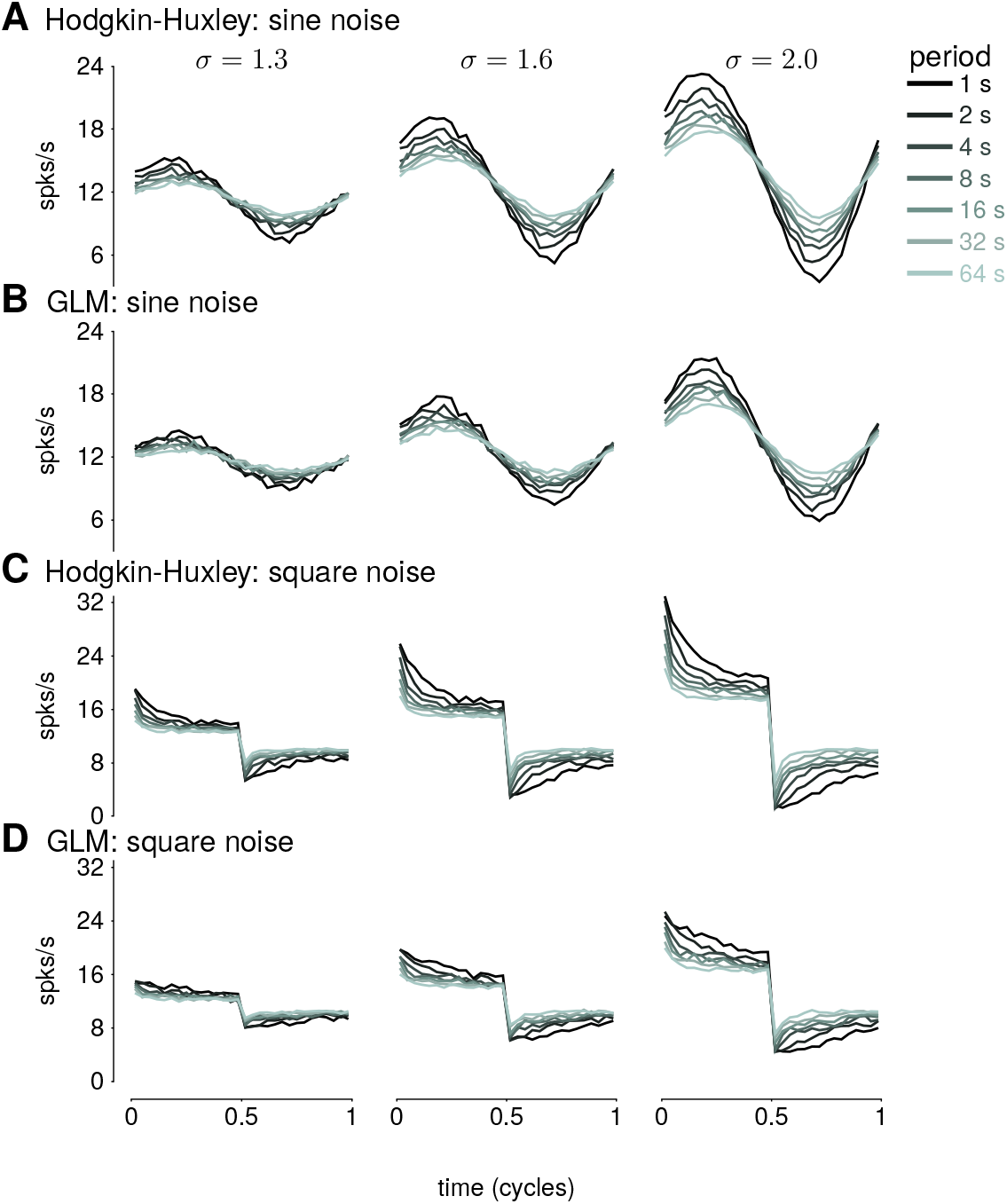
(**A**) The cycle-averaged response of the simulated Hodgkin-Huxley neurons with three AHP currents to sine wave modulated noise. Each trace shows the average response for a different period of noise modulation. The columns show the responses to different strengths of stimulus noise modulation (*σ*). (**B**) The cycle-averaged response of a GLM fit to the HH simulations in A. The GLM used a 16 s spike history filter. (**C**) The cycle-averaged response of the HH neurons to square-wave modulated noise. (**D**) The cycle-averaged response of a GLM fit to the HH simulations in C. The cycle averages can be compared to the exact fractional derivatives in **Figure 2B,D**.

The sinusoidal noise simulations show two properties of fractional differentiation. First, we estimated response gain (i.e., the strength of the sinusoidal modulation in the cycle-averaged response as a function of stimulus period; **Figure 9A**). In an ideal fractional differentiator, the log gain is proportional to the log of the stimulus period. The HH neuron shows a near linear response (*r*^2^ = 0.99, *p* – 10^−4^). Although the GLM with short history shows an almost flat relationship, increasing the spike history length shows similar slope to the HH neuron. The second property was the phase lead of the cycle-averaged response relative to the stimulus (**Figure 9B**). The phase lead should be constant under perfect fractional differentiation. The phase lead declines with longer period, but the HH simulation still shows strong phase lead in a 64 s period. Short spike history filter GLMs exhibit a phase lead that tends to zero with long SD periods. However, the GLM fit with a long spike history filter closely tracks the HH neuron’s phase lead.

**Figure 9:**
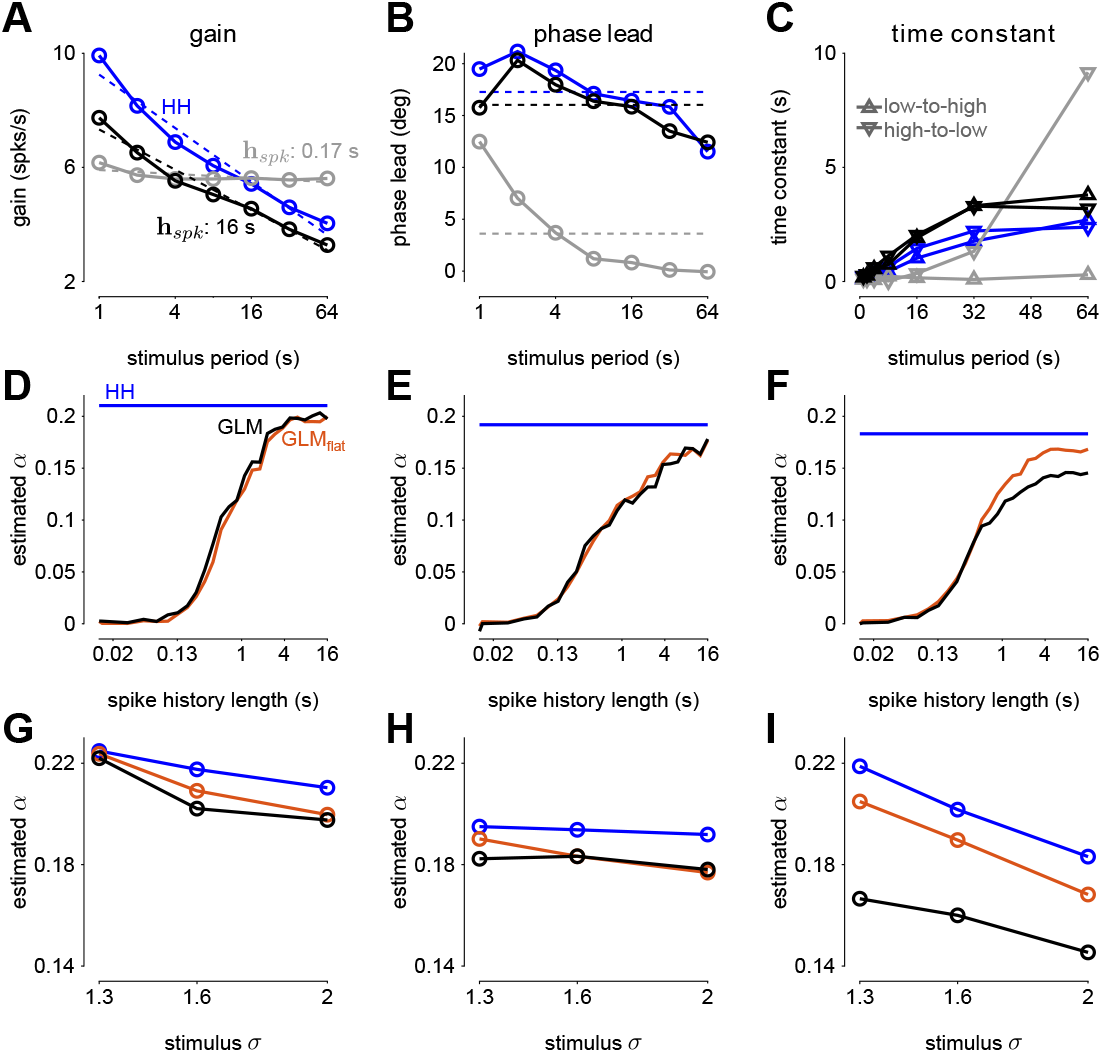
(**A**) The gain of the average responses to sine-wave modulated noise as a function of stimulus period for the HH and GLMs (**Figure 8A-B right**). The GLM fit with a 0.17 s spike history filter (gray) is compared to the GLM with the full 16 s spike history (black). The HH simulation is given in blue. The noise stimulus was modulated with *σ* = 2. (**B**) The phase lead of the average responses to sine-wave modulated noise as a function of stimulus period for the HH and GLMs. The fraction of variance explained in the HH phase lead curve by the GLM with 16 s spike history was *R*^2^ = 0.61. (**C**) The time constant of an exponential function fit to the cycle-averaged response to square wave noise for each stimulus period (**Figure 8C-D right**). The markers denote time constants estimated for steps from low to high variance or step from high v to low. The fraction of variance explained of the log time constants of the HH simulation by the GLM with 16 s spike history was *R*^2^ = 0.80. (**D**) The fractional differentiation order (*α*) of the GLM estimated by the slope of gain as a function of the log stimulus period in B. The value is estimated for each spike history lengths (black) and compared to *α* estimated from the HH simulation (blue). The red trace shows *α* estimated from the GLM fit only to unmodulated noise. (**E**) *α* estimated by the average phase lead across stimulus periods. (**F**) *α* estimated by fitting a the square-wave responses with a fractional differentiating filter. (**G-I**) *α* estimated at different noise modulation strengths for the 16 s spike history GLM and HH simulation.

The final signature of fractional differentiation was the exponential decay of the cycle-averaged response under square-wave noise simulation (**Figure 9C**). We estimate the time constant of the decay on the square noise cycle average for both steps up and steps down in stimulus SD. The time constant increases approximately linearly with the SD period, and GLMs with long spike history showed time constants closely approximated the HH neuron.

From each signature, we estimated the order of the fractional differentiation (*α*) in both the HH neurons and the GLM fits. We estimated the order using the slope of log-period compared to log-gain and mean phase lead across all stimulus periods for the sine-wave SD simulations (**Figure 9D-E**). A least-squares fit of FD filter of order *α* was applied to the square noise stimuli (**Figure 9F**). We considered *α* for the GLM fits as a function of the spike history length. The order estimates for the HH neuron, although slightly different for each signature, were approximately *α* = 0.2. The GLM’s FD order tends toward that of the HH neuron as the spike history length increases from below. Surprisingly, when we considered a GLM trained only to a flat noise stimulus (no sine or square modulation; stimulus SD *σ* = 1.0) showed similar *α* estimates (**Figure 9D-F**, red traces). Thus, the response properties giving rise to fractional differentiation of the noise envelope could be detected by the GLM even without driving with long timescale noise modulation.

We then considered how the estimated fractional differentiation order depended on the strength of the SD modulation. We found a slightly higher *α* for lower stimulus SDs. (note that *σ* = 2.0 was used to fit the GLMs) for the gain and timescale estimates (**Figure 9G-I**). However, the phase lead estimate was fairly stable across SDs.

Next, we quantified how well the GLM predicted the HH responses to new stimuli. Long timescale spike history filters improved the GLM’s ability to predict spike trains, and the improvement continued for spike histories of several seconds (**Figure 10A**). However, training only on unmodulated noise did not result in a good GLM fit despite predicting *α* (**Figure 10A**).

**Figure 10:**
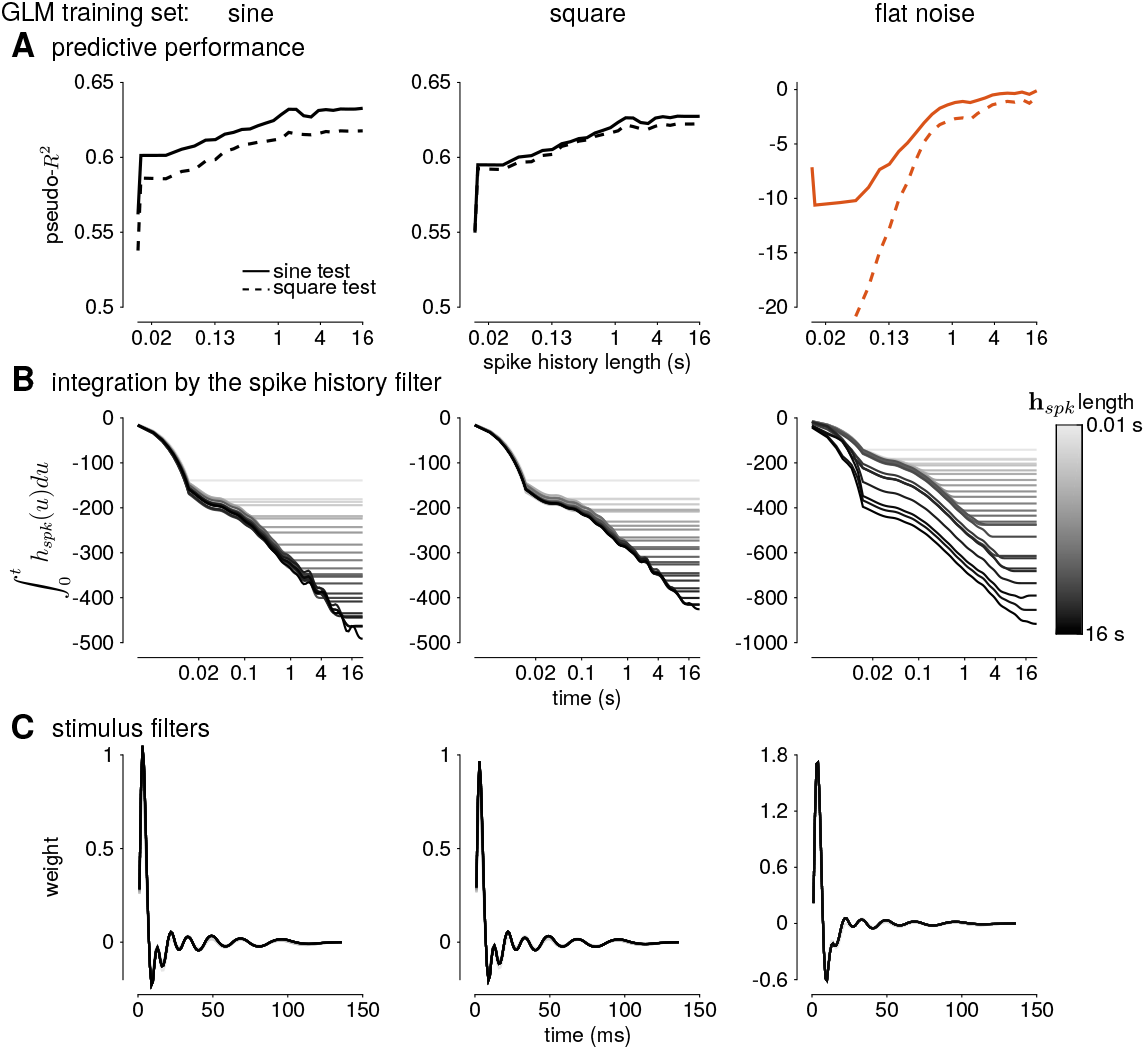
Assessing the model fitness for GLMs fit to sine wave modulated noise (left), square wave noise (middle), and unmodulated noise (right). (**A**) The pseudo-*R*^2^ measured on a withheld training set simulated from the HH model as a function of spike history length. The stimulus was sine (solid lines) or square (dashed lines) modulated noise with a 4 s period and a modulation strength of *σ* = 2. (**B**) The integral over time (i.e., cumulative sums) of each spike history filter. (**C**) The stimulus filters for all GLM fits.

We examined the parameter estimates in the GLM as a function of spike history length. We plotted the integral of the spike history filter to show how the filter integrates spikes over time. The integrals show long timescales seen for the GLM fit to either sine- or square-wave noise (**Figure 10B**). The GLM fit to either type of noise predicted over 60% of the variance in the HH responses to both sine- and square-wave noise. The flat noise GLM also showed long timescales, but the integral changed substantially with the spike history length changes. This indicates that the combination of spike-history dependent timescales is not well-constrained in the flat noise condition despite predicting *α*, perhaps due to biases present in the data without modulations (Stevenson, 2018). The stimulus filters are short timescale and showed little dependence on spike history length (**Figure 10C**). Thus, the GLM captured fractional differentiation in the HH neuron by linearizing the long timescale AHP currents.

## 4 Discussion

Individual neurons can adapt their responses to changes in input statistics. Here, we studied two adaptive computations to changes in the stimulus variance that are captured by biophysically realistic neurons. First, we examined gain scaling of the inputs so that the spike-triggered stimulus distribution was independent of the stimulus variance. The ability of the neuron to gain scale depended on the ratio of the spike-generating potassium and sodium conductances. Second, we considered spiking responses that approximate a fractional derivative of the stimulus standard deviation, which can be produced by a set of AHP currents with different timescales. Although HH neurons can produce these adaptive effects, it is difficult to fit the HH to data.

Our results demonstrate that the GLM provides a tractable statistical framework for modeling adaptation to stimulus variance in single-neurons. The GLM provides an alternative representation of the spiking responses as two linear filters (stimulus and spike history filters) with a fixed spiking nonlinearity instead of a multidimensional (and potentially stochastic) dynamical system (Meng et al., 2011, 2014). Importantly, a single GLM could accurately approximate the responses of HH neurons across multiple levels of input variance or across multiple timescales of variance modulation. The GLM accomplished this by linearizing the effect of recent spiking into a nonlinear and stochastic spiking mechanism to adjust for the current stimulus statistics. To reproduce gain scaling, only around 150 ms of spike history is required, in line with the rapid expression of the gain scaling property with changes in stimulus statistics (Fairhall et al., 2001a; Mease et al., 2013). In the fractional derivative case, the GLM summarized the multiple AHP currents of the HH models as a single linear autoregressive function with long timescale effects.

The simulations explored here assumed the input to a cell was an injected current generated from a Gaussian distribution. However, neurons receive input as excitatory and inhibitory conductances, which can be integrated across complex dendritic processes. Additionally, realistic input statistics may not follow a Gaussian distribution. Further work towards understanding the adaptive computations performed by single neurons should consider the inputs the neuron receives within a broader network.

Neural coding and computations that occur across a wide range of input levels depend heavily on adaption to the stimulus variance (Wark et al., 2007). The GLM, despite being a simple approximation, can provide a good representation of adaptive computations in biophysically realistic neurons.

## Author Contributions

KL and AF designed the study. KL performed the simulations and statistical analysis. KL and AF wrote the the manuscript.

## Acknowledgments

I We thank Jonathan Pillow and Misha Proskurin for their help in developing this project.

## Data Availability Statement

All simulations are publicly available at https://github.com/latimerk/GainScalingGLM.

## Supplementary Information

**Supplementary Figure 1:**
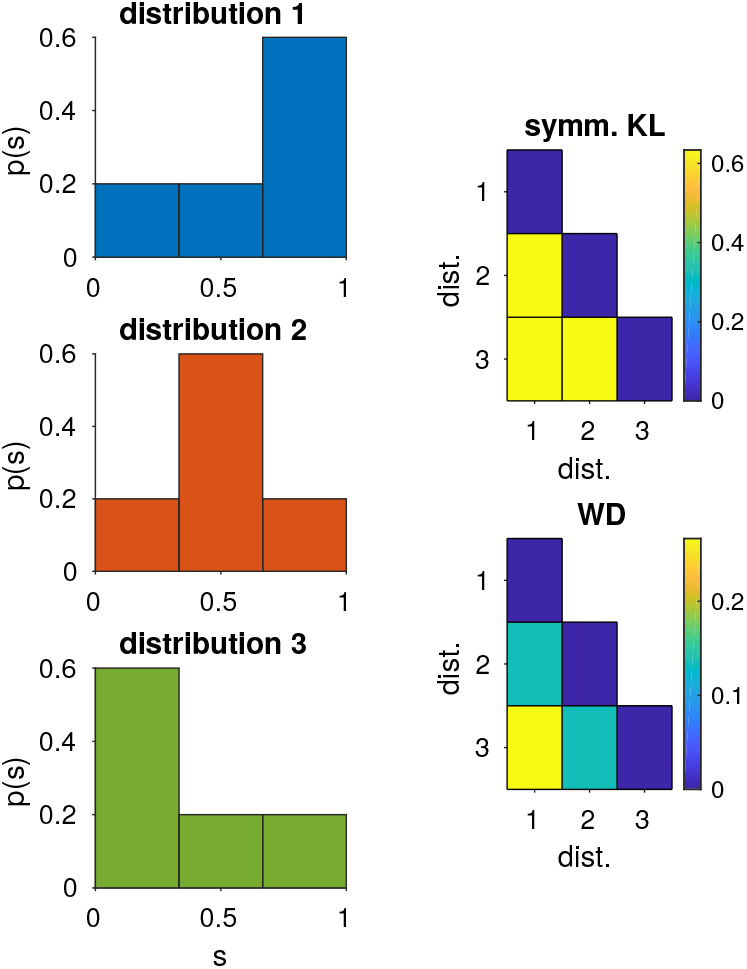
Example differences between the Kullback-Leibler (KL) divergence with the Wasserstein distance used to measure gain scaling. The difference in the three distributions is probability mass moved along the axis. (Left) Three example distributions over the value *s* are given. (Right) The (symmetrized) KL divergence between each of the three distributions and the Wasserstein distances. The Wasserstein metric depends on the distance the peak of probability mass is moved along the axis: the distance between distributions 1 and 3 is greater than between distributions 1 and 2. In contrast, the KL divergence does not depend on the distance and the divergence between each pair of distributions is equal.

**Supplementary Figure 2:**
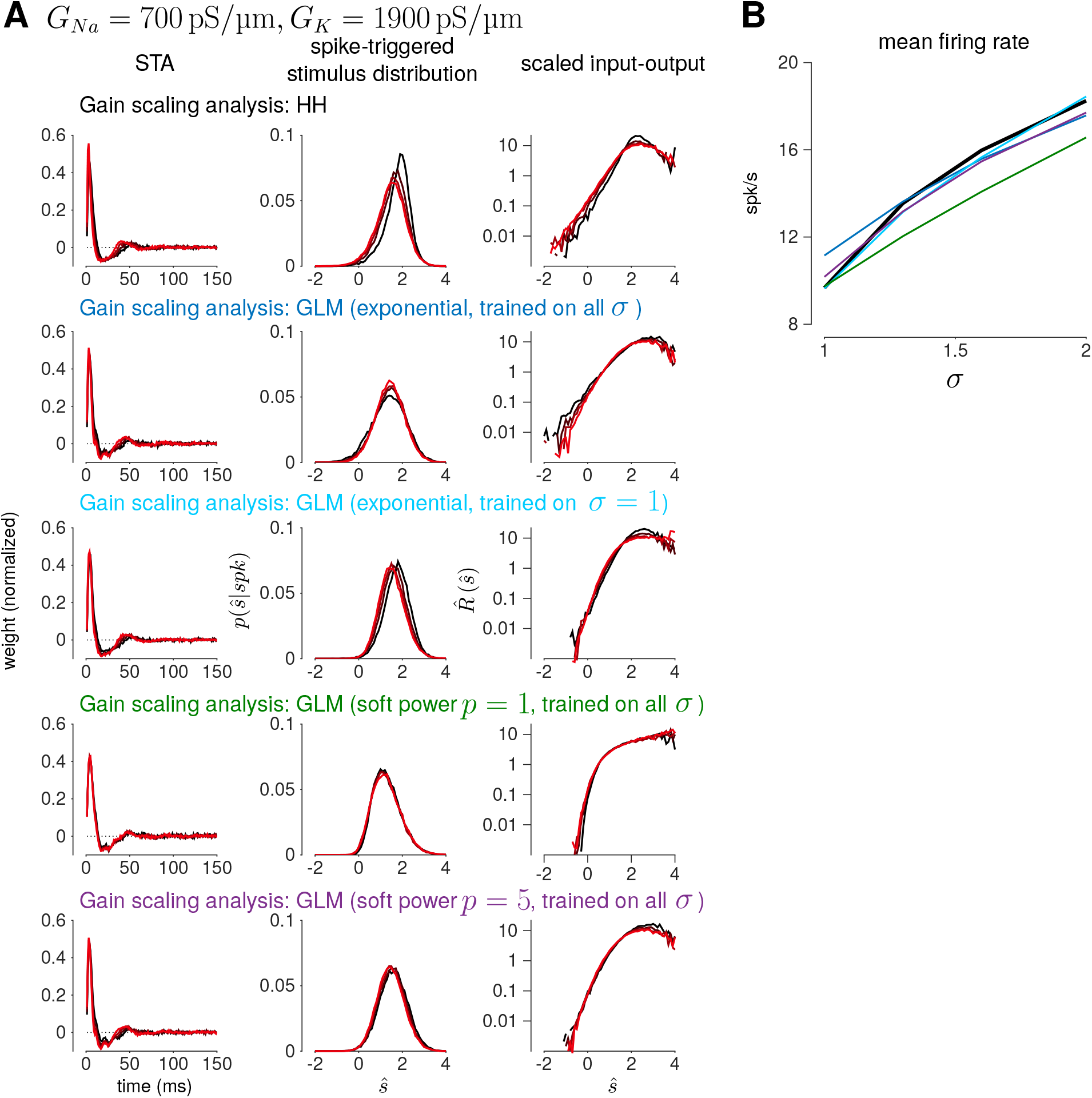
(**A**) Example gain scaling analysis for four different GLM fits to the HH simulation in the top row. The HH simulation has a low sodium to potassium ratio with poor gain scaling. (**B**) The firing rates of the HH and GLM simulations as a function of *σ*.

**Supplementary Figure 3:**
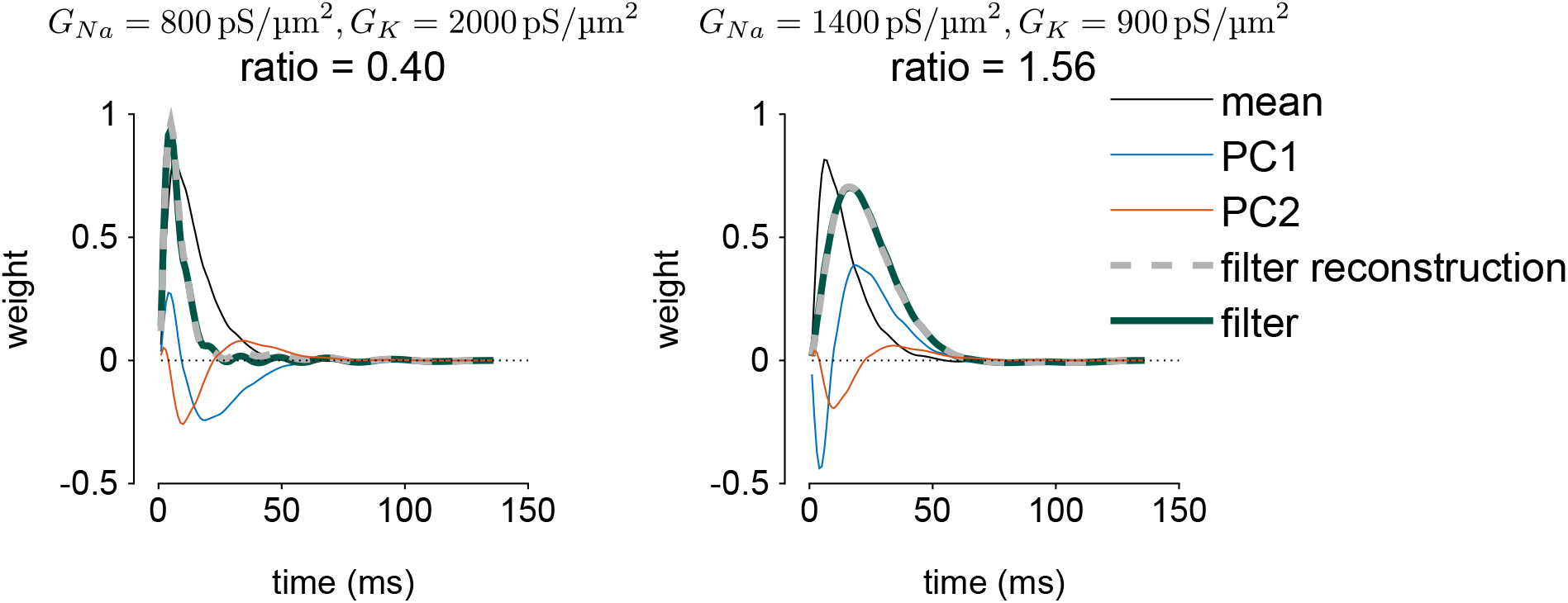
Example PCA reconstruction of the GLM’s stimulus filter for two of the HH fits. The mean filter and the two weighted PC vectors are given. The filter is reconstructed from the 2-D PCA space as the sum of the mean and the two PCs (dashed gray trace), and the reconstruction can be compared to the GLM filter (dark teal trace). Adding the weighted combinations of the two PCs extends or shortens the mean filter instead of adding multiple modes.

